# A Common Representational Code for Event and Object Concepts in the Brain

**DOI:** 10.1101/2025.08.22.671793

**Authors:** Jia-Qing Tong, Jeffrey R. Binder, Lisa L. Conant, Stephen Mazurchuk, Andrew J. Anderson, Leonardo Fernandino

**Author notes:** Corresponding Author: Leonardo Fernandino, Departments of Neurology and Biomedical Engineering Medical College of Wisconsin, 8701 W Watertown Plank Rd, Milwaukee, WI 53226. Financial interests or conflicts of interest: None.

## Abstract

Events and objects are two fundamental ways in which humans conceptualize their experience of the world. Despite the significance of this distinction for human cognition, it remains unclear whether the neural representations of object and event concepts are categorically distinct or, instead, can be explained in terms of a shared representational code. We investigated this question by analyzing fMRI data acquired from human participants (males and females) while they rated their familiarity with the meanings of individual words (all nouns) denoting object and event concepts. Multivoxel pattern analyses indicated that both categories of lexical concepts are represented in overlapping fashion throughout the association cortex, even in the areas that showed the strongest selectivity for one or the other type in univariate contrasts. Crucially, in these areas, a feature-based model trained on neural responses to individual event concepts successfully decoded object concepts from their corresponding activation patterns (and vice versa), showing that these two categories share a common representational code. This code was effectively modeled by a set of experiential feature ratings, which also accounted for the mean activation differences between these two categories. These results indicate that neuroanatomical dissociations between events and objects emerge from quantitative differences in the cortical distribution of more fundamental features of experience. Characterizing this representational code is an important step in the development of theory-driven brain-computer interface technologies capable of decoding conceptual content directly from brain activity.

**Significance Statement:** We investigated how word meaning is encoded in the brain by examining the neural representations of individual lexical concepts from two distinct categories—objects and events. We found that both kinds of concepts were encoded in neural activity patterns in terms of a shared representational space characterized by different modalities of perceptual and emotional experience. This indicates that individual concepts from a wide variety of semantic categories can, at least in principle, be decoded from neural activity using a generative model of concept representation based on interpretable semantic features. Furthermore, both object and event concepts could be decoded from cortical regions previously hypothesized to encode category-specific representations, suggesting that the two categories are jointly represented in these areas.

## Introduction

The nature of the distinction between object concepts (e.g., *bottle*, *dog*, *mountain*) and event concepts (e.g., *storm*, *party*, *hiccup*) has long been debated among philosophers and cognitive scientists (Casati, 2020; Zacks and Tversky, 2001). Prototypical objects have properties that remain constant over time, such as shape, size and mass, and have definite spatial borders. Typically, two objects cannot co-exist in the same location. Events, on the other hand, are necessarily tied to the notion of time, typically having beginnings and endings. Unlike objects, events are not mutually exclusive with respect to spatial location and do not have clear spatial borders.

From a neuroscientific perspective, object and event concepts appear to map onto partly distinct sets of cortical areas, as indicated by neuropsychological studies of individuals with brain injury (Aggujaro et al., 2006; Caramazza and Hillis, 1991; Damasio and Tranel, 1993; Daniele et al., 1994; McCarthy and Warrington, 1985; Miceli et al., 1988; Miceli et al., 1984; Shapiro and Caramazza, 2003) and by functional neuroimaging experiments (Bedny et al., 2014; Davis et al., 2004; Elli et al., 2019). Specifically, object concepts have been associated mainly with cortical areas along the ventral temporal cortex, overlapping the visual “what” pathway (Milner and Goodale, 2008; Mishkin et al., 1983), whereas event concepts are more strongly associated with the posterior lateral temporal cortex (Bedny et al., 2014; Kable et al., 2002; Kemmerer et al., 2008). This neuroanatomical dissociation has been interpreted as evidence that object and event concepts are encoded as categorically distinct types of representations rather than relying on a shared representational system (Bedny et al., 2008; Bedny and Caramazza, 2011).

The present study investigated whether this dissociation can be accounted for with a feature-based model of concept representation in which a common code underlies both kinds of concepts (**Figure 1**). We evaluated this hypothesis by testing five empirical predictions: both object-favoring and event-favoring cortical areas (defined as those areas that are more strongly activated by one of these concept types relative to the other) encode information about concepts in the non-favored category (P1); event and object concepts can be decoded from neural activations based on the same featural encoding model (P2); neural activation patterns for individual concepts in one category can be decoded by a featural encoding model trained exclusively on data from the other category (P3); the differential activation to these two concept categories, both in event-favoring and object-favoring areas, can be predicted by the tuning profiles to individual model features exhibited by these areas (P4); and finally, in event-favoring areas, the difference in activation between categories is better predicted by the features that are most uniquely important for characterizing events (i.e., “event-salient” features such as *harm*, *sound*, *consequential*); conversely, in object-favoring areas, the difference in activation is better predicted by “object-salient” features (e.g., *motion*, *taste*, *manipulability*) (P5).

**Figure 1.**
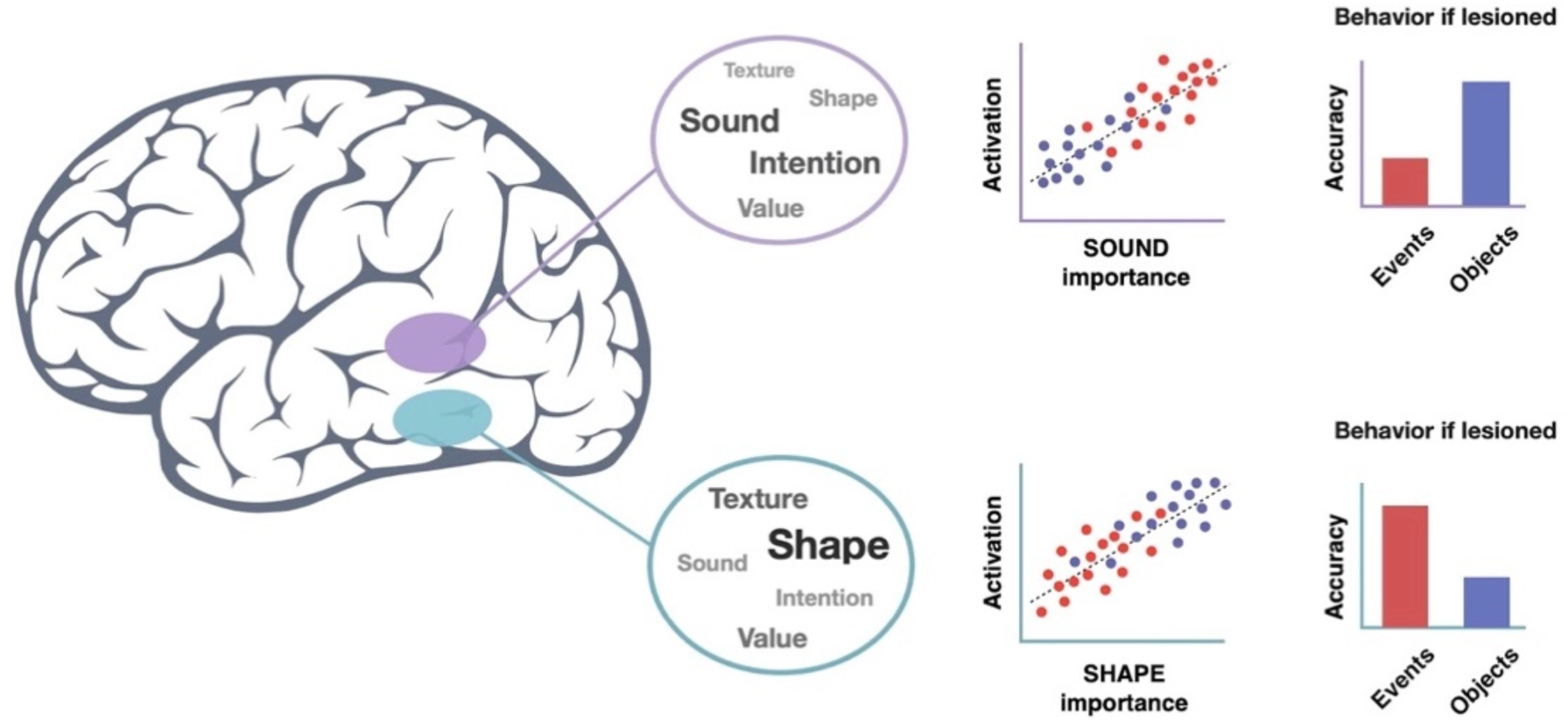
Schematic example of how a shared representational code could lead to category-specific effects in functional neuroimaging and brain damage studies. Although the hypothetical purple and green regions represent the same types of information, the purple region represents more of the information that is critical for recognizing and knowing about events, whereas the green region represents more of the information that is critical for recognizing and knowing about objects.

We first used univariate contrasts to identify cortical areas that are more active when processing concepts from one category relative to the other, thus determining the event-favoring and the object-favoring networks. Then, using a previously validated set of experiential semantic features based on distinct dimensions of experience (Binder et al., 2016), we conducted representational similarity analysis (RSA) on multi-voxel activation patterns to test whether each of these networks encodes information about individual concepts from both categories. We then used multivariate machine-learning techniques (voxelwise encoding models) to determine whether activation to individual object concepts could be decoded based on event concept data alone, and vice versa. Feature importance analyses identified subsets of experiential features that were most uniquely important for discriminating among concepts in each category (henceforth, event-salient and object-salient features; **Table 1**), and encoding model analyses investigated which feature subset best predicted neural activity in each network. Finally, we assessed whether the univariate contrast between events and objects could be predicted by the responsiveness profile of each point in the cortical sheet with respect to experiential features.

**Table 1.** Event- and object-salient feature sets (top-performing features for classifying event and object concepts into subcategories).

|  | *Feature* | *Description* |
| --- | --- | --- |
| **Event-Salient** | Cognition | Something that is a form of mental activity or a function of the mind |
|  | Harm | Someone or something that could cause harm to you or others |
|  | Communication | A thing or action that people use to communicate |
|  | Social | An activity or event that involves an interaction between people |
|  | Complexity | Something that is visually complex |
|  | Benefit | Someone or something that could help or benefit you or others |
|  | Sound | Having a characteristic or recognizable sound or sounds |
|  | Speech | Someone or something that talks |
|  | Fearful | Someone or something that makes you feel afraid |
|  | Consequential | Likely to have consequences (cause other things to happen) |
|  | Large | Large in size |
|  | Vision | Something that you can easily see |
|  | Unpleasant | Someone or something that you find unpleasant |
| **Object-Salient** | Biomotion | Showing movement like that of a living thing |
|  | Path | Showing changes in location along a particular direction or path |
|  | Motion | Showing a lot of visually observable movement |
|  | Fast | Showing visible movement that is fast |
|  | Taste | Having a characteristic or defining taste |
|  | Smell | Having a characteristic or defining smell or smells |
|  | Head | Associated with actions using the face, mouth, or tongue |
|  | Human | Having human or human-like intentions, plans, or goals |
|  | Body | Having human or human-like body parts |
|  | Upper Limb | Associated with actions using the arm, hand, or fingers |
|  | Away | Associated with movement away from or out of you |
|  | Manipulation | A physical object you have personal experience using |
|  | Color | Having a characteristic or defining color |

## Materials and Methods

The fMRI data set, briefly described below, was originally reported in Study 2 of Fernandino et al. (2022), to which we refer the reader for a more extensive description.

### Participants

Participants were 39 healthy, right-handed, native English speakers (21 women, 18 men; mean age, 28.7), with no history of neurologic disease. Participants were compensated for their time and gave informed consent in conformity with a protocol approved by the Institutional Review Board of the Medical College of Wisconsin.

### Stimuli

The stimulus set consisted of 160 object nouns (40 each of animals, plants/foods, tools, and vehicles) and 160 event nouns (40 each of social events, verbal events, non-verbal sound events, and negative events; **Table 2**). Object and event concepts were matched on letter and phoneme length, number of syllables, orthographic bigram statistics, orthographic and phonologic neighborhood density, word frequency, and mean naming and lexical decision reaction time (Balota et al., 2007).

**Table 2.** Stimulus items in the experiment.

| Event |  |  |  | Object |  |  |  |
| --- | --- | --- | --- | --- | --- | --- | --- |
| Negative | Nonverbal Sound | Social | Communication | Animal | Food/Plant | Tool | Vehicle |
| avalanche | applause | banquet | advice | alligator | asparagus | anchor | ambulance |
| battle | bang | bash | apology | ant | banana | axe | automobile |
| blizzard | bellowing | carnival | class | baboon | bean | baseball | barge |
| bombing | boom | celebration | commemoration | bison | beer | binoculars | bicycle |
| brawl | chattering | christening | comment | butterfly | blueberry | book | boat |
| cyclone | chuckle | circus | commentary | cardinal | bread | calculator | bobsled |
| downpour | clapping | cocktails | complaint | caterpillar | broccoli | camera | bus |
| drought | clattering | concert | compliment | chameleon | carrot | candle | canoe |
| earthquake | crackle | conference | debate | cheetah | champagne | cash | car |
| epidemic | crescendo | contest | denial | chicken | cheese | comb | carriage |
| explosion | giggle | convention | deposition | chimpanzee | cherry | corkscrew | convertible |
| famine | groaning | cookout | dictation | chipmunk | chestnut | crutches | elevator |
| flood | growling | cruise | discourse | cricket | chocolate | dime | escalator |
| gunshot | grunt | dance | dispute | crow | cider | faucet | ferry |
| gust | gulp | expedition | eulogy | dog | coffee | football | glider |
| hail | hiccup | expo | greeting | dolphin | cranberry | fork | helicopter |
| hailstorm | jingle | fair | grievance | duck | cucumber | glass | jeep |
| hurricane | laughter | feast | huddle | elephant | custard | hairbrush | limousine |
| inferno | melody | festival | interrogation | fish | dandelion | hammer | locomotive |
| invasion | murmuring | fiesta | joke | goldfish | egg | handsaw | motorcycle |
| landslide | reverberation | gathering | lecture | hamster | eggplant | hoe | plane |
| lightning | roaring | housewarming | lesson | hawk | flower | key | rocket |
| monsoon | rumble | jubilee | meeting | hippopotamus | ham | keyboard | rowboat |
| murder | rustle | luncheon | plea | horse | honey | ladle | sailboat |
| outbreak | screaming | march | praise | jackal | jam | magazine | scooter |
| plague | screeching | musical | protest | lion | ketchup | microscope | skateboard |
| raid | shrieking | outing | quarrel | monkey | lemonade | newspaper | sled |
| riot | sigh | pageant | rant | moose | milk | pencil | sleigh |
| shooting | siren | parade | rebuke | mosquito | mushroom | rake | steamer |
| squall | sizzle | party | rebuttal | mouse | mustard | sandpaper | streetcar |
| stampede | snap | picnic | recitation | octopus | nectarine | scissors | submarine |
| storm | sneeze | prom | sermon | penguin | pineapple | skillet | subway |
| tempest | sobbing | rally | showdown | rhinoceros | plant | spatula | taxi |
| thunderstorm | squeaking | reception | squabble | salmon | pudding | stapler | tractor |
| tornado | squeal | reunion | testimony | snake | pumpkin | stethoscope | train |
| twister | thumping | safari | thanks | tiger | raspberry | straw | tricycle |
| volcano | thunderclap | symphony | threat | trout | sauerkraut | thermometer | trolley |
| war | wheezing | tour | trial | turkey | spaghetti | ticket | truck |
| whirlwind | whimpering | tournament | tribute | turtle | tobacco | tongs | van |
| wildfire | whine | wedding | wisecrack | whale | tomato | umbrella | wagon |

### Task

Words were presented visually in a fast event-related design with a variable inter-stimulus interval. The entire list was presented to each participant six times in different pseudorandom orders across three separate imaging sessions (two presentations per session) on separate days.

On each trial, a noun was displayed for 500 ms, followed by a 2.5 s blank screen. Each trial was followed by a central fixation cross with variable duration between 1 and 3 s (mean = 1.5 s). Participants rated each noun according to how familiar they were with the corresponding entity or event on a scale from 1 (low) to 3 (high), indicating their response by pressing one of three buttons on a response pad with their right hand.

### MRI Data Acquisition and Preprocessing

Images were acquired with a 3T GE Premier scanner at the Medical College of Wisconsin Center for Imaging Research. Structural imaging included a T1-weighted MPRAGE volume (FOV = 256 mm, 222 axial slices, voxel size = 0.8 x 0.8 x 0.8 mm^3^) and a T2-weighted CUBE acquisition (FOV = 256 mm, 222 sagittal slices, voxel size = 0.8 x 0.8 x 0.8 mm^3^). T2*-weighted gradient-echo echoplanar images were obtained for functional imaging using a simultaneous multi-slice sequence (SMS factor = 4, TR = 1500 ms, TE = 33 ms, flip angle = 50°, FOV = 208 mm, 72 axial slices, in-plane matrix = 104 x 104, voxel size = 2 x 2 x 2 mm^3^). A pair of T2-weighted spin echo echo-planar scans (5 volumes each) with opposing phase-encoding directions were acquired before run 1, between runs 4 and 5, and after run 8, to provide estimates of EPI geometric distortion in the phase-encoding direction. Data preprocessing was performed using fMRIPrep 20.1.0 software (Esteban et al., 2019).

After slice timing correction, functional images were corrected for geometric distortion using nonlinear transformations estimated from the paired T2-weighted spin echo images. All images were then aligned to correct for head motion before aligning to the T1-weighted anatomic image. All voxels were normalized to have a mean of 100 and a range of 0 to 200. To optimize alignment between participants and to constrain the searchlight analysis to cortical gray matter, individual brain surface models were constructed from T1-weighted and T2-weighted anatomic data. We visually checked the quality of reconstructed surfaces before carrying out the analysis. Segmentation errors were corrected manually, and the corrected images were fed back to the pipeline to produce the final surfaces. The cortex ribbon was reconstructed in standard grayordinate space with 2 mm spaced vertices, and the EPI images were projected onto this space. For the following univariate contrast analysis, the EPI images were smoothed with a 6 mm FWHM kernel.

### Experiential Semantic Features

The experiential model of word meaning is based on the 65-dimensional space described by Binder et al. (Binder et al., 2016). In brief, the feature set was designed to cover the entire spectrum of human phenomenological experience from a cognitive neuroscience perspective, based in part on known distinctions between the neural processing systems underlying sensory-motor, affective, and social experience. Volunteers rated the relevance of each experiential feature to a given lexical concept on a 7-point Likert scale. This feature set is highly effective at clustering concepts into canonical taxonomic categories and has been used successfully to model semantic information encoded in fMRI activation patterns during word and sentence reading (Anderson et al., 2017; Anderson et al., 2019; Fernandino et al., 2022; Tong et al., 2022). Previous comparisons with other lexical semantic models, including WordNet, word2vec, GloVe, and Small World of Words showed that experiential feature ratings are better at predicting the similarity structure of activation patterns throughout the association cortex, and that these other models do not add predictive power beyond that achieved by experiential feature ratings (Fernandino et al., 2022; Tong et al., 2022; Tong and Fernandino, 2024).

### Feature Importance Analysis

The set of event concepts and the set of object concepts were each composed of items belonging to 4 subcategories, with 40 items in each subcategory. We estimated the relative importance of each experiential feature for the representation of event and object concepts, separately, by assessing the extent to which a feature contributes to the classification of these concepts into these subcategories. We used a composite score of three different metrics: two univariate filter methods – analysis of variance (ANOVA) and mutual information – and a multivariate wrapper method, namely, random forest classification with recursive feature elimination and cross-validation (RF-RFECV). ANOVA determined the extent to which a feature explained between-subcategory variance relative to within-subcategory variance, assuming that features that explain the most between-subcategory variance contribute the most to categorization. The mutual information score between each feature rating and the categorical variable denoting the *a priori* subcategories provided a measure of feature importance that, unlike ANOVA, does not assume linear dependency between the two variables.

In RF-RFECV, the least important feature was removed after each classifier training/testing iteration until a single feature remained. The order in which each feature was removed was used as the ranking of importance. For the random forest estimator, the performance was tested based on the out-of-bag accuracy score. There were 5000 trees per iteration, and all features were used to build each tree. The analysis was implemented using Python Scikit-learn (Pedregosa et al., 2011). Separately for object and event concepts, the features were ranked according to each of these methods, and a composite result of feature importance was produced by averaging the three rankings. Rankings produced by the three methods were compared using pair-wise correlations (Spearman’s Rho).

For each concept type, we selected the top k most important features, with k being the largest subset size for which the two subsets of top-performing features had no features in common, resulting in 13 object-salient features and 13 event-salient features. These features were used in the analysis “Network Encoding Model with Event- and Object-Salient Features” below.

### Univariate Contrast Analysis

A general linear model was used to fit the time series of the fMRI data via multivariable regression. In the model, events and objects were each treated as a single regressor of interest and convolved with a hemodynamic response function, resulting in 2 beta coefficient maps for each participant. Regressors of no interest included head motion parameters (12 regressors) and response time (z-scores). Other nuisance regressors included word form-related variables, namely, number of letters, number of phonemes, number of syllables, word frequency, bigram frequency, trigram frequency, orthographic neighborhood density, phonological neighborhood density, mean phonotactic probability for single phonemes, and mean phonotactic probability for phoneme pairs, all obtained from the South Carolina Psycholinguistic Metabase (https://www.sc.edu/study/colleges_schools/artsandsciences/psychology/research_clinical_facilities/scope). We also included z-scored concreteness ratings (Brysbaert et al., 2014) and z-scored familiarity ratings. The familiarity rating for each word was obtained by averaging the participant’s responses across the 6 trials in which it appeared. The beta coefficients for events and objects were used to define category-favoring areas for subsequent analyses.

Category-favoring areas were defined by a two-tailed paired t-test comparing the beta coefficients for events and objects at each vertex on the cortical surface. Permutation Analysis of Linear Models (PALM; (Winkler et al., 2014) was used for nonparametric permutation testing to determine cluster-level statistical inference (10,000 permutations). We used a cluster-forming threshold of z > 3.1 (p < 0.001) and a cluster-level significance level of α < 0.01. The final data were rendered on the group-averaged HCP template surface. The event-favoring network was defined as the vertices showing significantly stronger activation to events, while the object-favoring network was defined as the vertices showing significantly stronger activation to objects.

### Word Map Generation

A general linear model was used to fit the EPI time series via multivariable regression. Each word (with its six presentations) was treated as a single regressor of interest. Regressors of no interest included 6 degrees of freedom of head motion, their temporal derivatives, and response time (z-scores) for the task. Regressors were convolved with a hemodynamic response function. The resulting 320 beta-coefficient maps were used in subsequent vertex-wise encoding analyses. For RSA, we used noise-normalized values (t maps), following the recommendation of Walther et al. (Walther et al., 2016).

### Representational Similarity Analysis

We used RSA to test whether each network represents conceptual content for the non-favored category—that is, whether the event-favoring network represents information about object concepts and whether the object-favoring network represents information about event concepts (prediction P1). For each network in each participant, we extracted a vector from each word activation map consisting of the values in each of the vertices in the network. We then computed the Pearson correlation between the vectors for each pair of words in a given category, thus generating a neural RDM for each category and for each network. The experiential feature ratings were used to generate separate model-based RDMs for object concepts and for event concepts. To account for the effects of potential confounding factors on the RSA, 13 control RDMs were constructed—one based on a categorical model coding for the four subcategories in each category, one based on concreteness ratings, one based on the subject’s own familiarity ratings, and one for each of the 10 word form-related variables mentioned above. RDMs for individual variables were calculated based on the Euclidean distance. Ordinary least squares regression was used to regress the control RDMs from each model-based RDMs. After the regression, Spearman correlations between each residual model RDM and each neural RDM were computed to obtain the final RSA scores. As an estimate of the total explainable variance in the neural RDMs, the lower bound of the noise ceiling was computed for each network as the mean, across participants, of the Spearman correlation between each participant’s neural RDM and the mean neural RDM averaged across all other participants.

A one-tailed one-sample t-test was performed for each correlation score across all participants. A two-way repeated ANOVA was performed across participants to investigate the interaction between Network and Category. A *post hoc* analysis was performed to compare the mean difference between the scores in each network. All *p*-values were corrected for multiple comparisons with FDR correction.

### Overview of the Vertex-Wise Encoding Model Analyses

All remaining analyses used encoding-decoding models based on ridge regression to evaluate whether, and to what extent, activations for individual concepts in a given set could be decoded based on a particular ensemble of experiential feature ratings, separately for each vertex. For a subset of the words (the training set), z-scored beta values were used as the dependent variable, with the corresponding z-scored experiential feature ratings as independent variables, resulting in a β coefficient for each feature at each vertex. To account for the subcategorical structure of the stimulus set (i.e., the four subcategories within each category), we adopted a controlled training approach; instead of splitting the data between training and testing sets in random fashion, we trained each model on data from one subcategory (e.g., animals) and tested its ability to predict data from other subcategories (e.g., tools). This approach enabled us to evaluate the ability of the experiential featural model to decode semantic representations based on differences among individual concepts rather than based on categorical distinctions. The ridge regression penalty term was determined as the one that produced the best average prediction score in a bootstrap procedure with 50 repetitions, with 4-fold cross-validation (trained on 30 beta coefficients to predict 10 left-out beta coefficients), testing values ranging from 1 to 1,000 (log scale). The resulting feature coefficient maps were then used to predict the activation for each concept in the test set based on the feature ratings for that word.

To account for variance explained by potential confounding variables, we also trained a ridge regression model consisting of nuisance factors (i.e., word-form variables, concreteness, and familiarity) to predict the data in the training set. We then used an ordinary least squares regression model, with the predicted values as the independent variable and the observed data as the dependent variable, to obtain the residualized test set data. The encoding model consisting of the semantic feature ratings was evaluated by computing the Pearson correlation between observed and predicted values across all words in the residualized test set data. All p-values were corrected for multiple comparisons with FDR correction. We conducted four vertex-wise encoding-decoding analyses to evaluate predictions P2-P5, as described below.

### Within-Category Decoding with All 65 Experiential Features

To formally test whether the same set of experiential features can decode both event and object concepts (prediction P2), we trained a vertex-wise encoding model consisting of the 65 feature ratings on the fMRI data for a subset of the concepts and used it to predict the activation for each of the held-out concepts, separately for each category. Since each category has 4 subcategories, there were a total of 12 possible training and test set combinations within each category. The prediction accuracy scores were averaged across all 12 combinations to produce the final score for each vertex. The average in each network, computed as the mean across vertices, was tested for significance with a one-tailed one-sample t-test across participants.

### Cross-Category Decoding with All 65 Experiential Features

The previous analysis cannot rule out the possibility that each of the two categories relies exclusively on one of two informationally independent representational codes captured by distinct subsets of features (i.e., the possibility that event decoding could have relied on a representational code that carries no information about object representation and vice versa, with the two representational codes being captured separately by distinct, informationally independent subsets of features). To rule out this possibility (prediction P3), we tested whether a vertex-wise encoding model based on the same feature ratings and trained on the fMRI data for concepts in one category could decode neural activation patterns for concepts in the other category. Successful prediction would indicate that a common representational code underlies the representation of event and object concepts in that vertex. There were 16 possible combinations of training and testing sets in each encoding-decoding direction (e.g., trained with events to predict objects and vice versa). We averaged the accuracy scores across all combinations in each direction, generating two scores at each vertex. The mean scores in each network were obtained by averaging each score across all of its vertices. They were then tested for significance with a one-tailed, one-sample t-test against zero across participants.

### Within-Category and Cross-Category Decoding Using Category-Salient Features

The goal of these analyses was to test the prediction that the event-favoring and the object-favoring networks are differentially tuned to particular experiential features (P4). Here, the encoding models were based on either the event- or the object-salient features (see section *Feature Importance Analysis* above). For within-category decoding, models were trained on the beta values of the words in one subcategory and evaluated based on the beta values of the words in a different subcategory within the same category. To quantify the unique variance explained by the event-salient feature model, we first computed residualized beta values by regressing out the predictions of a composite model consisting of the object-salient features along with the nuisance variables (i.e., concreteness, familiarity, and the word form-related variables). We then calculated the correlation between the predictions of the event-salient feature model and the residualized beta values. An analogous procedure was conducted to assess the unique variance explained by the object-salient feature set. These procedures generated four scores at each vertex. The mean scores in each network were calculated by averaging the scores across vertices and tested for significance with a one-tailed one-sample t-test across participants. To test whether the feature tuning profiles were different in the two networks, the scores were entered into a three-way repeated-measures ANOVA with Network, Category, and Feature Set as factors. A two-way repeated-measures ANOVA was then performed for each category to explore the interaction between Network and Feature Set.

Cross-category decoding used event-salient and object-salient encoding models trained in one subcategory to predict activations for items in subcategories in the other category (e.g., training on activations for animals and predicting activations for social events). As in the previous analysis, we used residualized beta values to evaluate the unique explanatory value of each category-salient encoding model, accounting for the variance explained by the other set of category-salient features and potential confounding lexical variables. There were 16 possible combinations of training and testing sets in each direction.

Averaging the accuracy scores in each direction resulted in four scores at each vertex. Mean scores for each network were obtained by averaging the scores across its vertices and tested for significance with a one-tailed one-sample t-test against zero across participants. To assess whether the two networks were differentially tuned to individual features, the scores were entered into a repeated-measures ANOVA with Network, Training Direction, and Feature Set as factors. Because an interaction between Network and Feature Set was found, we performed a two-way ANOVA in each network.

### Prediction of Activation Differences in Each Network Using Category-Salient Features

The goal of this analysis was to investigate whether the overall difference in activation between event and object concepts observed in each network is predicted by the network’s tuning profile with respect to experiential features. For each of the two sets of category-salient features, an encoding model consisting of those features was trained on the beta values for the words in a given subcategory (e.g., animal) and used to predict the beta values for words in each of two other subcategories—one in the same category (e.g., tool) and one in the other category (e.g., social events). The predicted activations were z-scored, and the mean predicted activation difference between the two subcategories was then recorded for each vertex. The overall prediction accuracy in each network was assessed by computing the Pearson correlation between the predicted and observed activation differences across vertices. The variance explained by nuisance variables was accounted for using the same procedure as in the previous analyses. For each encoding-decoding direction, there were 48 possible combinations of training and test sets across the 16 subcategories. These 96 accuracy scores were averaged together, generating a score for each set of category-salient features in each network. The scores in each condition were entered into a one-tailed one-sample t-test against zero. A two-way repeated-measures ANOVA was used to test the interaction between Feature Set and Network. *Post hoc* analysis was performed to compare differences between the mean scores in each network.

### Whole-Cortex Prediction of Activation Differences at Each Vertex

Finally, we investigated the extent to which the whole-brain activation map for the contrast between events and objects (with no thresholding applied) could be predicted by the responsiveness of each vertex to different experiential features. We trained an encoding model at each vertex using ridge regression with z-scored beta values for the 160 event words as the dependent variable and the 65 experiential feature values for those words (also z-scored) as independent (i.e., predictor) variables, resulting in a whole-brain beta map for each experiential feature. Before the dependent variables were z-scored, the 10 control lexical variables, concreteness ratings, and familiarity ratings were regressed out from the dependent variable. The optimal ridge regression penalty term was identified as described in the previous sections. The beta weights computed for each feature were then used to predict the activation amplitude for each object concept relative to the mean activation for event concepts (because the event beta maps used to train the model were mean-centered). The predicted activation maps for all 160 object concepts were then averaged, resulting in a predicted activation map for the object word set relative to the event word set. The predicted activation map for events relative to the objects was computed using the same procedure, and the two maps were averaged at the individual participant level. The resulting maps were smoothed with a 6-mm FWHM Gaussian kernel, and a one-sample t-test at each vertex was implemented to generate a group-level predicted event-vs-object contrast map. We assessed the similarity between this map and the observed event-vs-object univariate contrast map by computing the vertex-by-vertex Spearman’s ρ between the two maps. Statistical significance was determined by permutating the feature labels in the encoding model to generate a null distribution (1,000 permutations). A result larger than the 95^th^ percentile of the null distribution was considered significant.

## Results

We first determined which regions in the cerebral cortex were reliably activated by events relative to objects (the event-favoring network) and vice versa (the object-favoring network), using categorical contrasts at each location (i.e., at each vertex in the cortical surface mesh; Figure 2). The event-favoring network consisted of regions in the left anterior superior temporal sulcus (peak in HCP area L_STSva), left posterior superior temporal sulcus/temporoparietal junction (peaks in areas L_PHT, L_PSL, and L_PGi), left inferior frontal gyrus (peak in area 45), left precuneus and posterior cingulate gyrus (peaks in areas L_POS2 and L_31PV), left retrosplenial cortex (peak in area L_RSC), and right occipital cortex (peaks in areas R_V8 and R_V4). Object-favoring areas were observed in the left posterior middle temporal gyrus extending into the inferior temporal sulcus (peak in area L_PH), and in the intraparietal sulcus (peak in area L_IP1). These networks were used as regions-of-interest (ROIs) in subsequent analyses.

**Figure 2.**
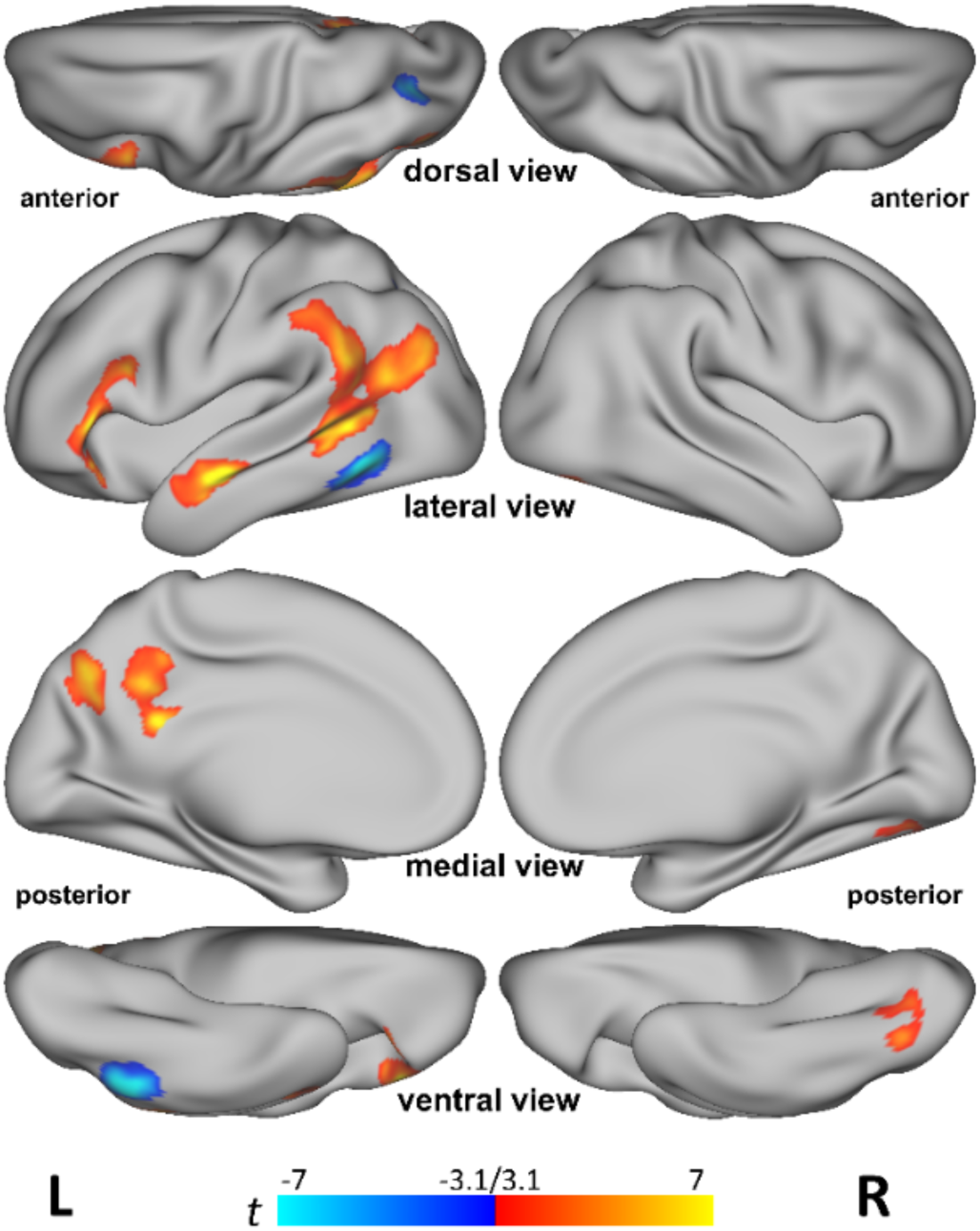
Univariate contrasts between event and object concepts. Red-yellow indicates stronger activation for event concepts and blue-cyan indicates stronger activation for object concepts.

### Event and object concepts are both represented in each of the two networks

As explained in *Materials and Methods*, in the multivariate analyses, the semantic content of each word was modeled as a vector of experiential feature ratings (Binder et al., 2016). As shown in Figure 3, objects and events differed reliably on many of these feature ratings and could be classified into *a priori* sub-categories with high accuracy based on the feature ratings alone.

**Figure 3.**
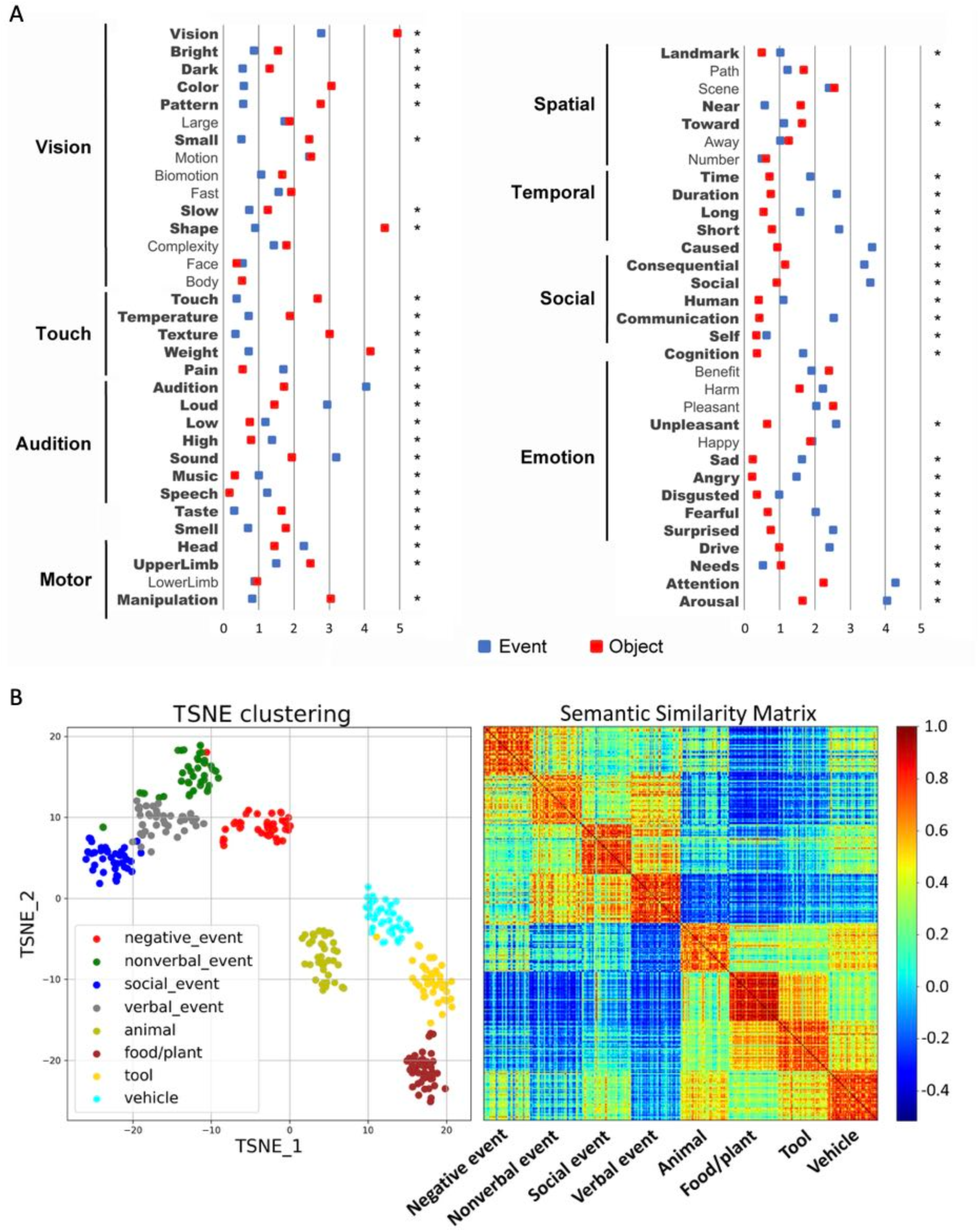
**A.** Mean feature ratings for the object (red) and event (blue) concepts. Bold font and * indicate significant difference (*p* < 0.05) after Bonferroni correction. **B**. T-distributed stochastic neighbor embedding (TSNE) clustering (left) and semantic similarity matrix (right) based on experiential features for 160 event concepts and 160 object concepts.

First, we investigated whether each network represented both event and object concepts by conducting RSAs based on this model (prediction P1). RSA provides a strong test of whether a brain region encodes a particular type of information about a set of stimuli. It assesses the degree to which differences between the neural responses to the stimuli reflect the differences predicted by the information encoded in the model-based RDM. In the present case, if the event-favoring network did not encode the experiential similarity structure of object concepts, similarities among neural responses to object nouns would show no correspondence to the similarities predicted by the semantic model, the same being true for neural responses to event stimuli in the object network.

As shown in Figure 4, all four RSA correlations were highly significant (event concepts in the event-favoring network: mean ρ = 0.077 ± 0.007, *p* < 0.001; object concepts in the object-favoring network: mean ρ = 0.032 ± 0.005, *p* < 0.001; object concepts in the event-favoring network: mean ρ = 0.033 ± 0.005, *p* < 0.001; event concepts in the object-favoring network: mean ρ = 0.043 ± 0.006, *p* < 0.001; all *p* values are FDR corrected). These results indicate that cortical areas that are differentially activated by event and object concepts are not dedicated exclusively to the representation of concepts in their favored category, as the semantic dissimilarity structures of both categories can be detected in each of the two networks. In the event-favoring network, the lower noise ceiling was 0.249 for events and 0.142 for objects. In the object-favoring network, the lower noise ceiling was 0.154 for events and 0.130 for objects. While there was a significant main effect of Category (F = 8.286, p = 0.006) showing overall higher RSA scores for event concepts, a significant interaction between Network and Category (F = 30.592, p < 0.001) shows that this advantage for event representation over object representation was restricted to the event-favoring network (t = 6.539, p < 0.001), with no difference between categories in the object-favoring network (t = −1.222, p = 0.229). This indicates that, despite the difference in mean activation observed in the univariate contrast, the two categories are similarly well-represented in the object-favoring network, while event concepts are substantially more strongly represented than object concepts in the event-favoring network.

**Figure 4.**
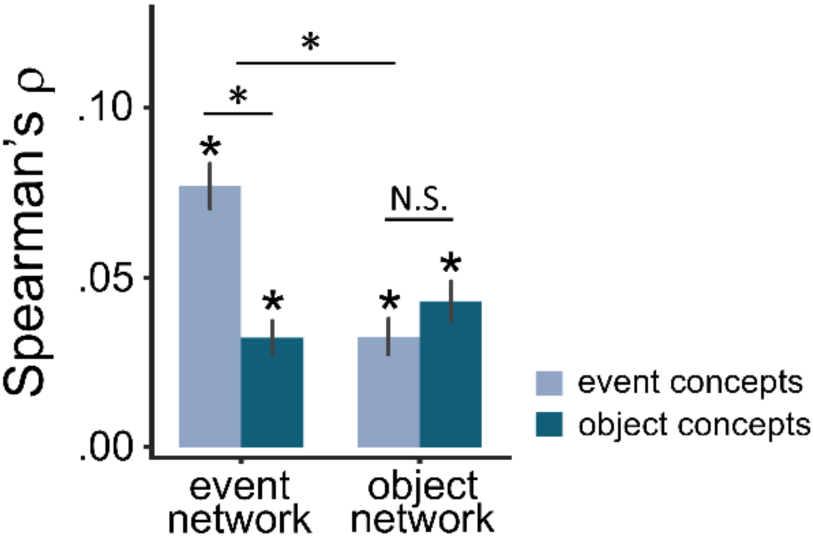
Results of the RSAs. Error bars represent the standard error of the mean. * *p* < 0.001.

### The same experiential model can decode both event and object concepts at the item level

The above-mentioned RSA results suggest that the same set of experiential features can potentially decode both event and object concepts. To formally test this prediction (P2), we trained a vertex-wise encoding model consisting of the 65 feature ratings on the fMRI data for concepts from one subcategory (e.g., tool) and used it to predict the activation for concepts from another subcategory (e.g., animal), separately for events and objects (see section *Vertex-Wise Encoding Model Analysis* in *Materials and Methods* for more details). The score was averaged across all vertices in each network, generating two scores per network per participant (i.e., training on events to decode events and training on objects to decode objects). The group-averaged correlation scores were significant in both networks (Figure 5B; all FDR-corrected p < 0.001), confirming that event and object representations can be decoded based on the same featural model (prediction #2), indicating that, in each category, a common representational code underlies different subcategories.

**Figure 5.**
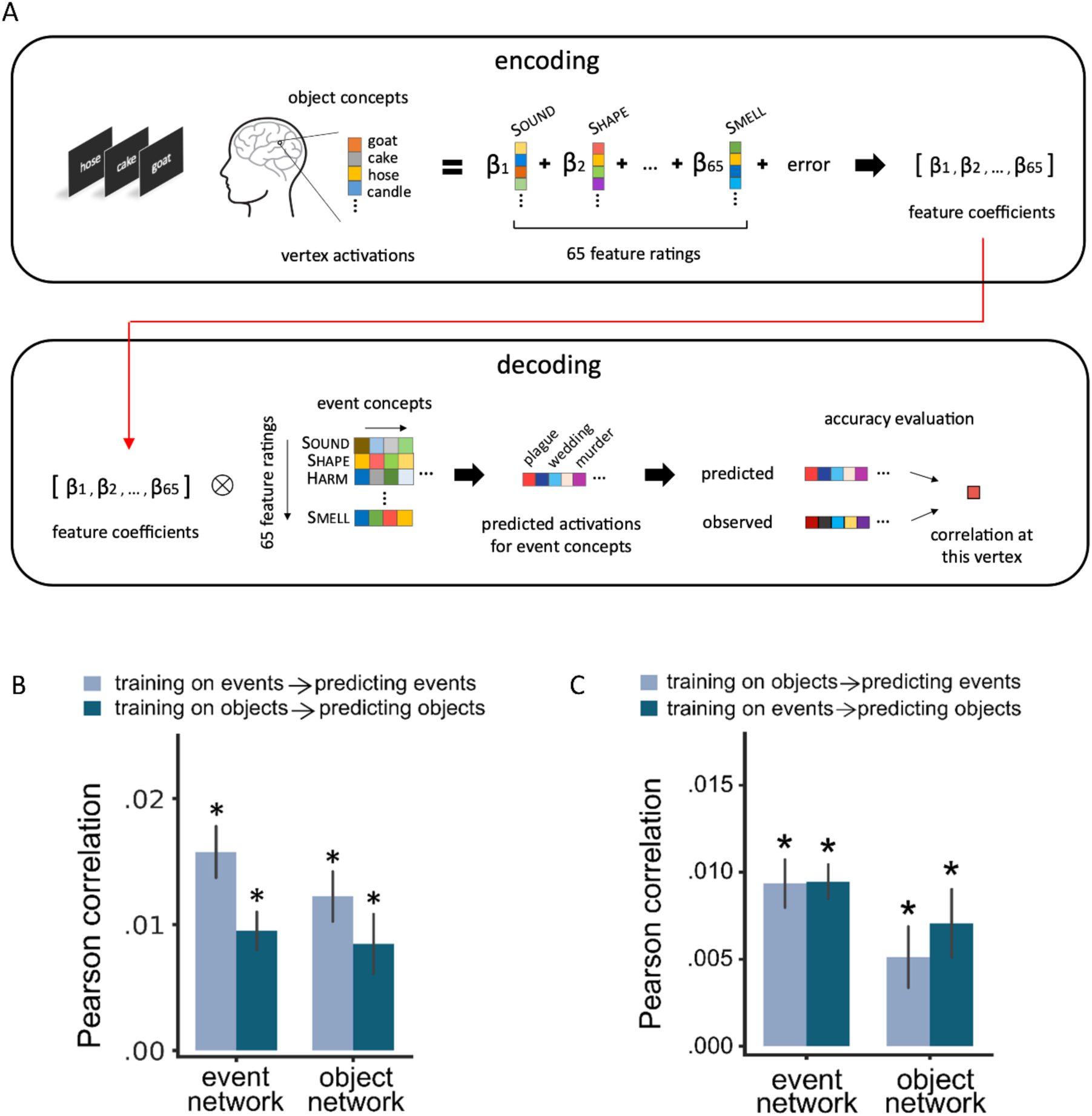
**A**. Schematic diagram of the encoding model of concept-specific activation. Prediction accuracy was evaluated via item-wise correlation with the observed values. **B**. Results of the within-category encoding model analysis, averaged across vertices in each network. Error bars represent the standard error of the mean. **C**. Results of the cross-category encoding model analysis, averaged across vertices in each network. Error bars represent the standard error of the mean. \**p* < 0.01.

### A common set of features accounts for the neural representation of both categories

The fact that concepts from both categories could be decoded based on the same set of 65 experiential features suggests that a single neural code may underlie the representation of both events and objects. To directly investigate this possibility, we tested whether a vertex-wise encoding model based on the same feature ratings and trained on the fMRI data for concepts in one category could decode neural activation for concepts in the other category (prediction P3; see section *Vertex-Wise Encoding Model Analysis* in *Materials and Methods* for more details). The score was averaged across all vertices in each network, generating two scores per network per participant (i.e., training on events to decode objects and training on objects to decode events). The group-averaged correlation scores were significant in both networks (Figure 5C; all FDR-corrected *p* < 0.001, except for models trained on objects to predict events in the object-favoring network: FDR-corrected *p* = 0.003), indicating that the two concept categories are represented in terms of a common code, and that this code can be modeled as a weighted combination of experiential feature ratings.

### Event- and object-favoring areas are differentially tuned to specific experiential features

Having shown that events and objects share a common representational code, we tested the prediction that the event-favoring and the object-favoring networks are differentially tuned to particular experiential features (P4). Specifically, we investigated whether the relative predictive power of two non-overlapping subsets of experiential features—namely, those that are most uniquely relevant to discriminating among subcategories within each category (see section *Feature Importance Analysis* in *Materials and Methods* for a detailed description of these analyses)—were different in the two networks. We expected that event-salient features would outperform object-salient features in the event-favoring network, and that the object-favoring network would show the opposite pattern. The top-ranking features for each category are listed in **Table 1**. At each vertex, activation data from a given subcategory was used to train a model based on either event- or object-salient features, and each model was used to generate predictions about the activation of individual concepts in a different subcategory within or across categories (see sections *Within-Category and Cross-Catego*ry *Decoding Using Category-Salient Features in Each Network* in *Materials and Methods* for more details).

In line with the previous analyses, the within-category encoding analysis showed that, in both networks, both event-salient and object-salient features explained unique variance in the neural activation to noun concepts, separately for events and objects (Figure 6C; all corrected *p* < 0.05). However, as predicted by the shared featural representation hypothesis (prediction P4), we observed a significant interaction between Network and Feature Set (Figure 6C; F = 24.525, p < 0.001), in which models consisting of event-salient features showed significantly higher accuracy than those consisting of object-salient features in the event-favoring network (F = 6.522, p = 0.015), while the object-favoring network showed the opposite pattern (F = 11.426, p = 0.002). No other interactions were found (all p > 0.7).

**Figure 6.**
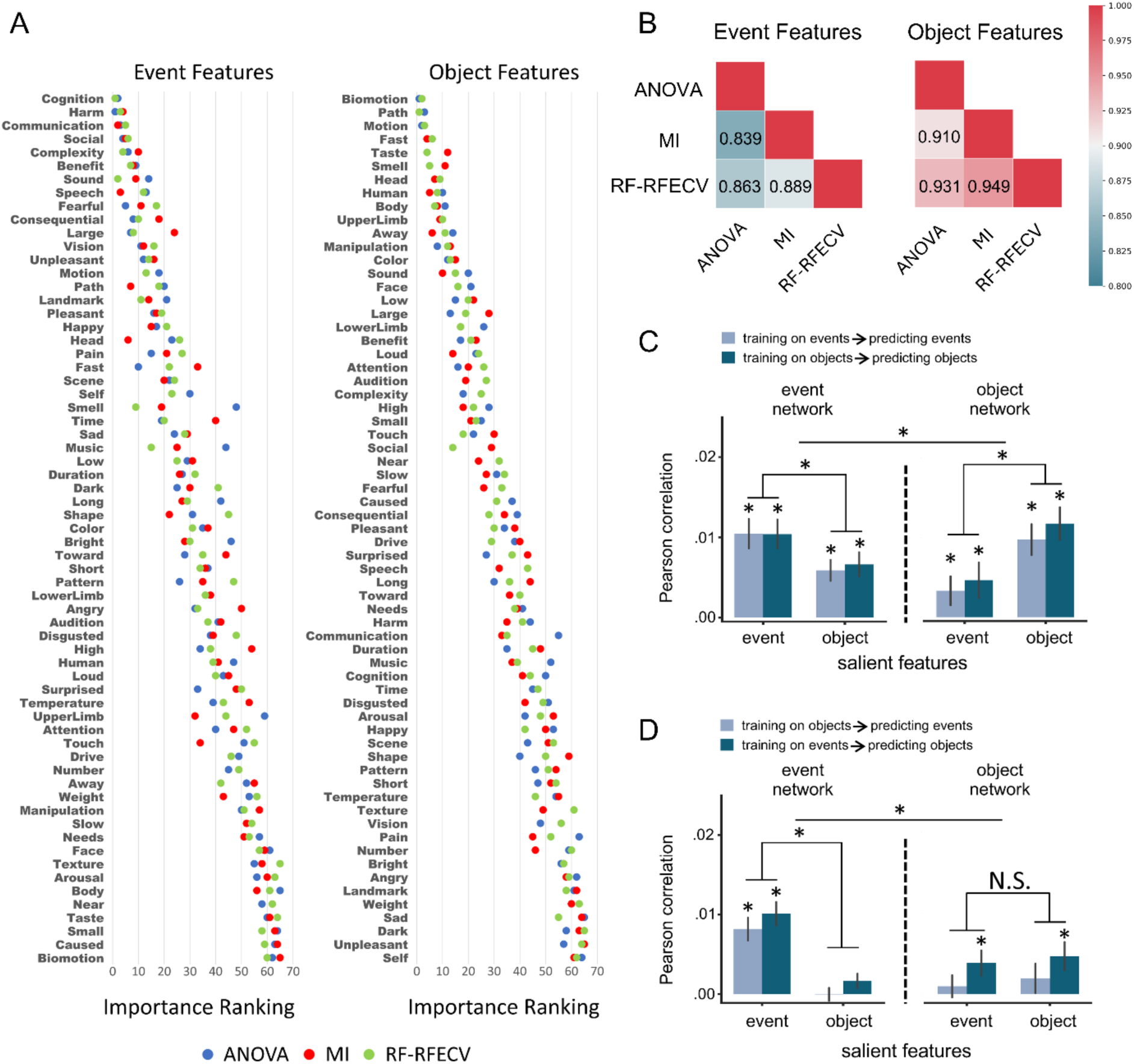
**A.** Feature importance for object and event concepts according to three different metrics. Rank order on y-axis was based on the mean of the three metrics. **B**. Pairwise correlations (Spearman’s rho) between ranks of feature importance generated according to each metric. ANOVA: analysis of variance; MI: mutual information; RF-RFECV: random forest with recursive feature elimination and cross-validation. **C**. Results of the within-category encoding model analysis based on category-salient features averaged across vertices in each network. **D**. Results of the cross-category encoding model analysis based on category-salient features averaged across vertices in each network. Error bars represent the standard error of the mean. \**p* < 0.05

Consistent with the within-category results, the cross-category encoding analysis also revealed a significant interaction between Network and Feature Set (F= 19.083, *p* < 0.001) in which models based on event-salient features outperformed those based on object-salient features in the event-favoring network (F = 24.091, *p* < 0.001) but not in the object-favoring network (F = 0.277, *p* = 0.602; Figure 6D). No other interactions were found (all *p* > 0.4).

Together, these findings indicate that, even though each network contributes to the neural representation of both categories, they preferentially encode information about features that are more salient to the category that produces the stronger mean activation in the univariate contrast. They suggest that, while information about each category is distributed throughout the association cortex, some regions contain relatively more information about objects or about events depending on how strongly tuned they are to different experiential features.

### The contrast between event and object concepts is predicted by the local tuning profile to experiential features

Finally, we investigated whether the differences in mean activation between event and object concepts observed in the univariate analysis could be predicted by the feature tuning profile in each region. The shared featural representation hypothesis predicts that the activation difference in the event-favoring network would be more strongly driven by event-salient features, whereas in the object-favoring network it would be more strongly driven by object-salient features (P5). We tested this prediction using a procedure similar to the one in the previous analysis, with encoding models based either on event-salient or object-salient features, separately for event-favoring and for object-favoring areas (Figure 7). We evaluated the accuracy of the predictions by computing the vertex-wise correlation between the observed and predicted activation differences between event and object concepts in each network (see section *Prediction of Activation Differences Using Salient Features* in *Materials and Methods* for more details).

**Figure 7.**
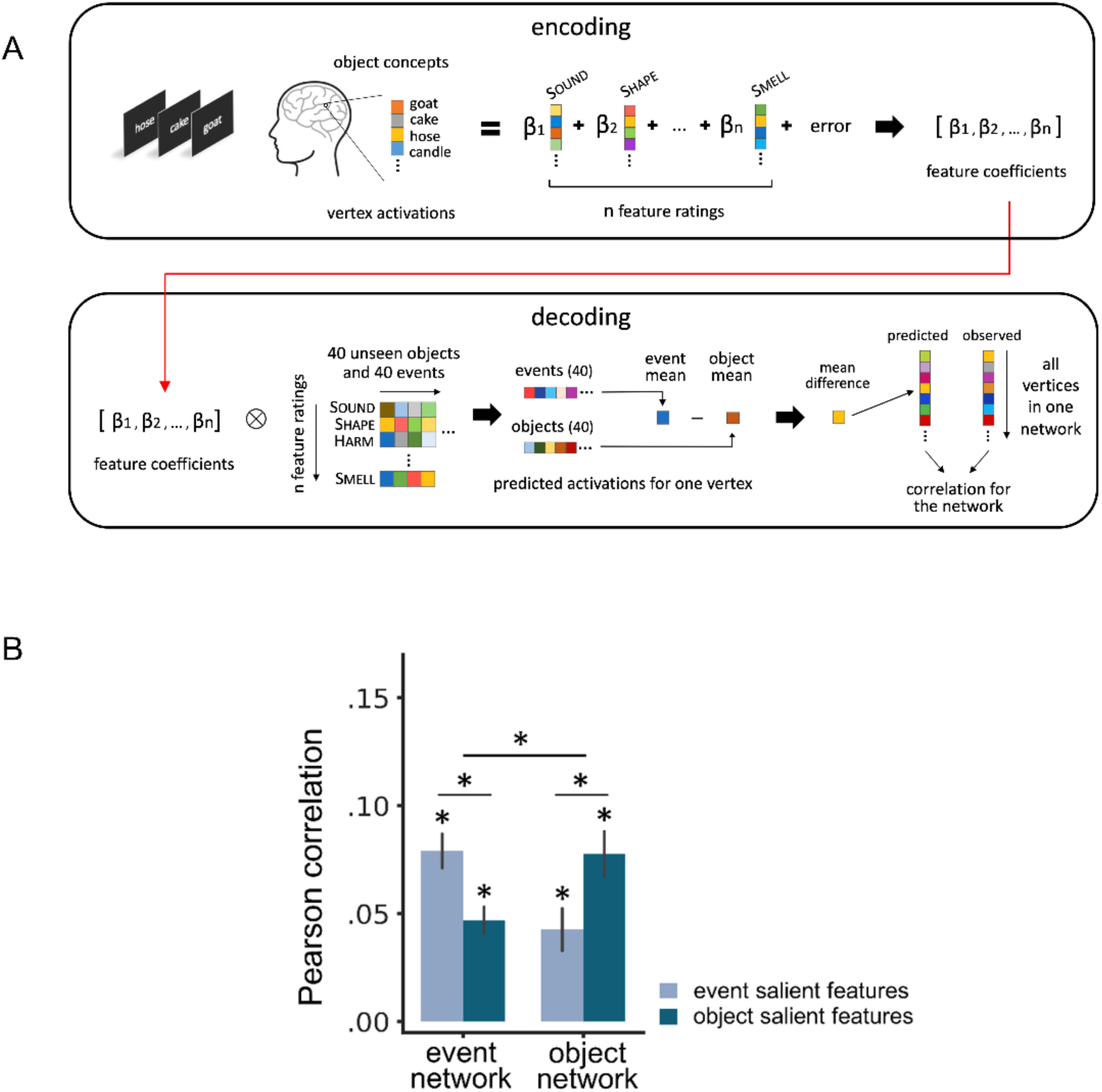
**A**. Schematic diagram of the encoding model of mean activation differences between objects and events. Prediction accuracy for each network was computed as the vertex-wise correlation with observed activations. **B**. Results of predicting the pattern of activation difference between events and objects using models based on category-salient features in each network. Error bars represent the standard error of the mean across participants; \**p* < 0.05.

As shown in Figure 7B, both models predicted the mean activation difference between categories significantly above chance in the event-favoring network (event-salient features: t = 9.909, FDR-corrected *p* < 0.001; object-salient features: t = 7.627, FDR-corrected *p* < 0.001) and in the object-favoring network (event-salient features: t = 4.301, FDR-corrected *p* < 0.001; object-salient features: t = 7.469, FDR-corrected *p* < 0.001). There was an interaction between Network and Feature Set (F = 16.406, FDR-corrected *p* < 0.001), such that models based on event-salient features had significantly higher scores than those based on object-salient features in the event-favoring network (t = 3.477, FDR-corrected *p* < 0.001), while the models based on object-salient features had significantly higher scores than those based on event-salient features in the object-favoring network (t = 2.576, FDR-corrected *p* = 0.014). These results indicate that activation differences between events and objects in univariate contrast analyses result from differences in the relative importance of individual experiential features across these categories allied to the fact that multimodal cortical areas are differentially tuned to particular experiential features.

While the analyses above were conducted on regions of interest determined by the univariate contrast between object and event concepts, our final analysis assessed whether the locations of those regions are predicted by the distributed experiential representation hypothesis. The difference map based on the local cortical responses to experiential feature ratings predicted the location of most category-favoring regions in the unthresholded contrast map, with a correlation across all cortical voxels that was highly significant (Spearman’s ρ = 0.373, *p* < 0.001; Figure 8).

**Figure 8.**
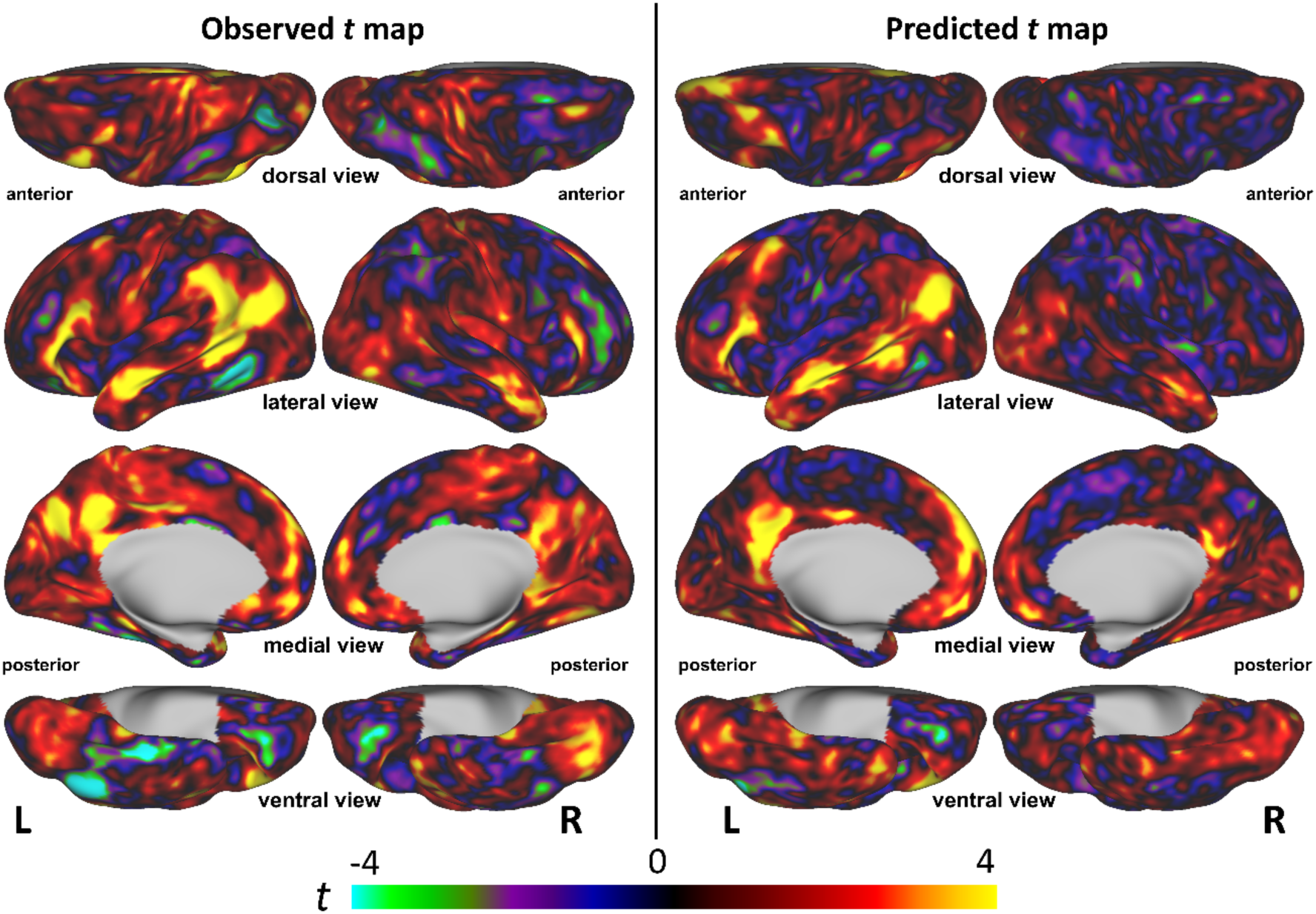
Results of the whole-cortex, vertex-wise encoding model based on all 65 experiential features. Warmer colors indicate stronger activation for event than for object concepts; cooler colors indicate stronger activation for object than for event concepts.

## Discussion

We investigated the proposal that lexical concepts referring to objects and events are jointly encoded in multimodal cortical areas according to a shared representational code that can be modeled with experiential feature ratings. We tested five key predictions of this hypothesis by applying machine learning techniques to fMRI data for individual lexical concepts and found that all five predictions were supported by the results.

The univariate contrast between event and object concepts identified cortical areas that activated significantly more strongly to one of these concept types (Figure 2). Because the two categories of word stimuli were matched in terms of grammatical category (i.e., all items were nouns), word frequency, and ten variables related to ortho-phonological word form, and because concept familiarity and concreteness were included as covariates, these potential confounding factors could be ruled out, teasing out neural activity related to conceptual content. The areas showing the strongest activations in the event-favoring network broadly match the results of similar contrasts in previous studies (Bedny et al., 2014; Elli et al., 2019; Martin et al., 1995) while highlighting the involvement of the left posterior cingulate gyrus and precuneus in event representation, which has not been previously reported. This novel finding is likely owed to the substantially higher statistical power of the present study allied to improvements in BOLD MRI acquisition and data preprocessing. In the only previous fMRI study that contrasted event and object nouns (Bedny et al., 2014), the peak temporal lobe activation for “events > objects” was in the inferior bank of the posterior superior temporal sulcus (pSTS; MNI coordinates −56, −48, 4), closely matching the present results. While that study reported another activation cluster in the left anterior middle and inferior temporal gyri, our results show, instead, a more dorsal cluster in the anterior STS. We also found that the left supramarginal gyrus was activated in this contrast, while in Bedny et al. this region was activated only for the contrast “verbs > nouns”.

The set of cortical areas showing stronger activation to objects (i.e., the object-favoring network) is similar to the network identified by the contrast “nouns > verbs” in Elli et al. (2019), in which nouns referred to either animals or places and verbs consisted of emission- and action-related concepts.

The multivariate fMRI analyses yielded converging evidence that event and object concepts are jointly represented in both sets of areas. RSAs of multivoxel activation patterns indicated that the regions showing the largest differences in mean activation between these two categories (i.e., the event-favoring and object-favoring areas) encoded semantic information about individual concepts in both categories, in agreement with prediction P1 (Figure 4). That is, in either set of areas, the similarity structure among distributed activation patterns for individual lexical concepts reflected the experiential similarity structure of both categories. In each network, the trend was toward more accurate decoding for the category showing the strongest mean activation, consistent with the idea that some areas are more critical to one category than to the other despite contributing to both. In conjunction with previous studies (Fernandino et al., 2022; Tong et al., 2022), these results suggest that all concepts are represented, to some degree, across the entire semantic network, such that no cortical region is exclusively dedicated to a single category.

The encoding-decoding analyses (Figure 5) indicate that lexical concepts from different superordinate categories are represented according to a shared representational code that can be successfully modeled by experiential feature ratings (prediction P2). Specifically, concepts in one subcategory (e.g., animals) could be decoded from neural activity using a feature-based model trained exclusively on data from a different subcategory (e.g., tools), which would be impossible if they relied on completely independent representational codes. Importantly, decoding was successful even when the encoding model was trained on objects and tested on events, and vice versa, in agreement with prediction P3.

While the experiential model captured shared variance across event and object concepts, it is relevant to consider the possibility that most of the 65 experiential features captured variance exclusively for one of the two categories, with only a few features capturing shared variance.

The results of the analysis *Cross-Category Decoding Using Category-Salient Features* argue against this possibility. This analysis identified the most uniquely important features for each category and then used linear regression to orthogonalize them with respect to each other. The finding that both residualized feature subsets were able to decode items in both categories shows that even the most uniquely distinctive experiential features of each category encoded information about the other category.

The encoding-decoding analyses using category-salient features showed that, in the event-favoring network, activation to individual concepts was more accurately predicted by event-salient features than by object-salient features, while concept representations in the object-favoring network were decoded more accurately by object-salient than by event-salient features (although in this network the difference did not reach significance in the cross-category analysis; Figure 6, panels C and D). This pattern of results indicates that the two networks have different tuning profiles to the experiential features examined (prediction P4, illustrated in Figure 1). This conclusion is further supported by our finding that, at event-favoring vertices, the difference in mean activation between object and event concepts is more accurately predicted by event-salient features, while at object-favoring vertices the difference is better predicted by object-salient features (prediction P5; Figure 7). The location of areas showing stronger activation for one category relative to the other could also be predicted from each voxel’s experiential tuning profile (Figure 8). Together, these findings show how category specialization in the brain can be seen as an emergent phenomenon arising from regional variation in experiential feature representation and systematic differences in experiential content between categories. They demonstrate that the neuroanatomical dissociations between objects and events reported in previous studies could be explained by parametric differences within a single representational system.

Is it possible that the results for event concepts in the object-favoring network were driven not by event representations but, instead, by object representations that are closely associated with those events (and vice versa)? Afterall, the fMRI activation pattern elicited in response to a single word (e.g., *famine*) may be affected by the automatic co-activation of thematically associated concepts in a different semantic category (*food*). However, thematic associations are not driven by experiential similarity; very dissimilar events such as *famine*, *cooking*, and *banquet* are each strongly associated with the object concepts *food* and *people*. Therefore, it seems unlikely that the neural similarity structure of the object associates of event concepts would substantially correlate with the experiential similarity structure of the original event concepts, or vice versa. The fact that events and objects exhibited RSA correlations of similar strength in the object-favoring network argues that activation of object associates is unlikely to completely account for the events RSA (Figure 4). The same rationale applies to the encoding-decoding analyses, particularly the ones based on category-salient features (Figure 6C). Nevertheless, this possibility cannot be entirely ruled out and should be investigated in future studies.

Linguistically, the categorical difference between events and objects is generally reflected in the grammatical distinction between verbs and nouns. As in most languages, English verbs typically refer to events (or event schemas) while most common nouns refer to temporally stable entities such as objects, people, places, and properties (Jackendoff, 2002). Although several functional neuroimaging studies have contrasted object nouns against verbs, we chose not to do so because of the unknown effects that grammatical class may have on the results.

However, as also noted by Elli et al. (2019), the observed differences in brain activity between nouns and verbs are similar to those observed between object and event nouns and are likely to be explained, to a large degree, by differences in experiential content. Notably, the areas activated in the contrast “events > objects” in the present study included the supramarginal gyrus, which was activated only in the “verbs > nouns” contrast in Bedny et al. (2014). These results provide additional evidence that activation in the “verbs > nouns” contrast reflects primarily semantic differences.

The model we used to represent experiential content has inherent limitations and is no doubt imperfect in its choice of features. It is an expanded version of previous similar models that represent content in terms of the relevance of particular ‘modes’ of experience to the acquisition of a concept (Fernandino et al., 2016; Gainotti et al., 2013; Hoffman and Lambon Ralph, 2013; Lynott and Connell, 2009, 2013). Some of these features are likely redundant due to covariances inherent in actual experience, and there likely are some important features missing. Nevertheless, the model outperforms a range of other models in explaining the similarity structure of word meanings as assessed via fMRI (Fernandino et al., 2022) and via automatic semantic priming (Fernandino and Conant, 2024). We speculate that one reason for this success may be that neural assemblies in these regions represent experience primarily in terms of the relative degree of activation of more modality-specific regions, rather than in terms of the detailed activation patterns that encode fine-grained information in each modality (Fernandino and Binder, 2024).

## Acknowledgements

This work was supported by National Institute on Deafness and Other Communication Disorders (NIDCD) grants R01 DC016622 and R01 DC020932, by the Intelligence Advanced Research Projects Activity under Grant FA8650-14-C-7357, and by a grant from the Advancing a Healthier Wisconsin Foundation (Project #5520462). The authors thank Volkan Arpinar, Elizabeth Awe, Joseph Heffernan, Steven Jankowski, Jedidiah Mathis, and Megan LeDoux for technical assistance.

## References

Aggujaro S, Crepaldi D, Pistarini C, Taricco M, Luzzatti C (2006) Neuro-anatomical correlates of impaired retrieval of verbs and nouns: Interaction of grammatical class, imageability and actionality. Journal of Neurolinguistics 19:175–194.

Anderson AJ, Binder JR, Fernandino L, Humphries CJ, Conant LL, Aguilar M, Wang X, Doko D, Raizada RDS (2017) Predicting neural activity patterns associated with sentences using a neurobiologically motivated model of semantic representation. Cereb Cortex 27:4379–4395.

Anderson AJ, Binder JR, Fernandino L, Humphries CJ, Conant LL, Raizada RDS, Lin F, Lalor EC (2019) An integrated neural decoder of linguistic and experiential meaning. J Neurosci 39:8969–8987.

Balota DA, Yap MJ, Cortese MJ, Hutchison KA, Kessler B, Loftis B, Neely JH, Nelson DL, Simpson GB, Treiman R (2007) The english lexicon project. Behav Res Methods 39:445–459.

Bedny M, Dravida S, Saxe R (2014) Shindigs, brunches, and rodeos: The neural basis of event words. Cogn Affect Behav Neurosci 14:891–901.

Binder JR, Conant LL, Humphries CJ, Fernandino L, Simons SB, Aguilar M, Desai RH (2016) Toward a brain-based componential semantic representation. Cogn Neuropsychol 33:130–174.

Brysbaert M, Warriner AB, Kuperman V (2014) Concreteness ratings for 40 thousand generally known english word lemmas. Behav Res Methods 46:904–911.

Caramazza A, Hillis AE (1991) Lexical organization of nouns and verbs in the brain. Nature 349:788–790.

Casati RaV, Achille (2020) Events. In: The Stanford encyclopedia of philosophy Summer 2020 ed Edition (Zalta EN, ed): Metaphysics Research Lab, Stanford University.

Damasio AR, Tranel D (1993) Nouns and verbs are retrieved with differently distributed neural systems. Proc Natl Acad Sci U S A 90:4957–4960.

Daniele A, Giustolisi L, Silveri MC, Colosimo C, Gainotti G (1994) Evidence for a possible neuroanatomical basis for lexical processing of nouns and verbs. Neuropsychologia 32:1325–1341.

Davis MH, Meunier F, Marslen-Wilson WD (2004) Neural responses to morphological, syntactic, and semantic properties of single words: An fMRI study. Brain Lang 89:439–449.

Elli GV, Lane C, Bedny M (2019) A double dissociation in sensitivity to verb and noun semantics across cortical networks. Cereb Cortex 29:4803–4817.

Esteban O, Markiewicz CJ, Blair RW, Moodie CA, Isik AI, Erramuzpe A, Kent JD, Goncalves M, DuPre E, Snyder M, Oya H, Ghosh SS, Wright J, Durnez J, Poldrack RA, Gorgolewski KJ (2019) Fmriprep: A robust preprocessing pipeline for functional MRI. Nat Methods 16:111–116.

Fernandino L, Binder JR, Desai RH, Pendl SL, Humphries CJ, Gross WL, Conant LL, Seidenberg MS (2016) Concept representation reflects multimodal abstraction: A framework for embodied semantics. Cereb Cortex 26:2018–2034.

Fernandino L, Tong JQ, Conant LL, Humphries CJ, Binder JR (2022) Decoding the information structure underlying the neural representation of concepts. Proc Natl Acad Sci U S A 119.

Fernandino L, Binder JR (2024) How does the “default mode” network contribute to semantic cognition? Brain and Language 252:105405.

Fernandino L, Conant LL (2024) The primacy of experience in language processing: Semantic priming is driven primarily by experiential similarity. Neuropsychologia:108939.

Gainotti G, Spinelli P, Scaricamazza E, Marra C (2013) The evaluation of sources of knowledge underlying different conceptual categories. Front Hum Neurosci 7:40.

Hoffman P, Lambon Ralph MA (2013) Shapes, scents and sounds: Quantifying the full multi-sensory basis of conceptual knowledge. Neuropsychologia 51:14–25.

Jackendoff R (2002) Foundations of language: Brain, meaning, grammar, evolution: Oxford University Press UK.

Kable JW, Lease-Spellmeyer J, Chatterjee A (2002) Neural substrates of action event knowledge. J Cogn Neurosci 14:795–805.

Kemmerer D, Castillo JG, Talavage T, Patterson S, Wiley C (2008) Neuroanatomical distribution of five semantic components of verbs: Evidence from fMRI. Brain Lang 107:16–43.

Lynott D, Connell L (2009) Modality exclusivity norms for 423 object properties. Behav Res Methods 41:558–564.

Lynott D, Connell L (2013) Modality exclusivity norms for 400 nouns: The relationship between perceptual experience and surface word form. Behav Res Methods 45:516–526.

Martin A, Haxby JV, Lalonde FM, Wiggs CL, Ungerleider LG (1995) Discrete cortical regions associated with knowledge of color and knowledge of action. Science 270:102–105.

McCarthy R, Warrington EK (1985) Category specificity in an agrammatic patient: The relative impairment of verb retrieval and comprehension. Neuropsychologia 23:709–727.

Miceli G, Silveri MC, Villa G, Caramazza A (1984) On the basis for the agrammatic’s difficulty in producing main verbs. Cortex 20:207–220.

Miceli G, Silveri MC, Nocentini U, Caramazza A (1988) Patterns of dissociation in comprehension and production of nouns and verbs. Aphasiology 2:351–358.

Pedregosa F, Varoquaux G, Gramfort A, Michel V, Thirion B, Grisel O, Blondel M, Prettenhofer P, Weiss R, Dubourg V, Vanderplas J, Passos A, Cournapeau D, Brucher M, Perrot M, Duchesnay E (2011) Scikit-learn: Machine learning in python. Journal of Machine Learning Research 12:2825--2830.

Shapiro K, Caramazza A (2003) Grammatical processing of nouns and verbs in left frontal cortex? Neuropsychologia 41:1189–1198.

Tong J, Binder JR, Humphries C, Mazurchuk S, Conant LL, Fernandino L (2022) A distributed network for multimodal experiential representation of concepts. J Neurosci 42:7121–7130.

Tong J-Q, Fernandino L (2024) Experiential information drives the neural decoding of word meaning in models based on word association. Highlights in the Language Sciences, Nijmegen, Netherlands.

Walther A, Nili H, Ejaz N, Alink A, Kriegeskorte N, Diedrichsen J (2016) Reliability of dissimilarity measures for multi-voxel pattern analysis. Neuroimage 137:188–200.

Winkler AM, Ridgway GR, Webster MA, Smith SM, Nichols TE (2014) Permutation inference for the general linear model. Neuroimage 92:381–397.

Zacks JM, Tversky B (2001) Event structure in perception and conception. Psychol Bull 127:3–21.

